# Medial prefrontal cortex encodes implicit temporal expectations in mice

**DOI:** 10.64898/2025.12.15.694406

**Authors:** Federica Lareno Faccini, Pierre Le Cabec, Ludovic Spaeth, Matthieu Pasquet, Dominique Ciocca, Anne Giersch, Philippe Isope

**Author notes:** Contributed equally (Federica Lareno Faccini, Pierre Le Cabec). Present address: Centre interdisciplinaire de recherche en biologie, Collège de France, CNRS, INSERM, 75005, Paris, France. Present address: Dominick P Purpura Department of Neuroscience, Albert Einstein College of Medicine, Bronx, New York, USA.

## Abstract

Temporal prediction allows animals to align their actions with upcoming events, yet most work has focused on explicit duration judgments rather than the implicit timing that shapes ongoing behavior. Here, we ask how medial prefrontal cortex implements such latent temporal expectations and how cerebellar input contributes. Head-fixed mice learned a cued water-delivery task in which reward occurred after either a short or long delay, or after a single fixed delay. During variable delays, running and licking became anticipatory, and medial prefrontal local field potentials and single neurons showed ramping and reward-locked activity patterns aligned to expected reward time. Switching to a fixed delay rapidly sharpened behavioral anticipation and temporal coding. Optogenetic activation of cerebellar Purkinje cells selectively perturbed these dynamics and biased behavior around the earliest possible reward time. These results identify a cerebello-prefrontal circuit that encodes implicit temporal predictions on the sub-second scale.

## Introduction

Temporal prediction is fundamental to everyday behavior, allowing animals and humans to align their actions with events unfolding over hundreds of milliseconds to seconds. Although the neural basis of explicit time estimation has been studied extensively, much less is known about the mechanisms that support the incidental (“implicit”) timing that shapes ongoing behavior in the absence of overt temporal judgements. Converging evidence from human neuroimaging and animal physiology points to distributed cortico–subcortical networks, including prefrontal cortex and cerebellum, as key substrates for representing temporal regularities and predicting the timing of expected outcomes (Ivry and Spencer, 2004; Coull and Nobre, 2008; Buonomano and Laje, 2011; Cravo et al., 2011; Merchant et al., 2013; Coull et al., 2016; Herbst and Obleser, 2019). However, how these regions cooperate to generate implicit temporal expectations, and how such expectations are expressed in specific population dynamics during naturalistic behavior, remains poorly understood.

The PFC is central to everyday skills—sequencing actions, holding goals in mind, staying flexible, and making decisions (Goldman-Rakic, 1996; Fuster, 2001; Miller and Cohen, 2001; Le Merre et al., 2021) — it is therefore well positioned to support the temporal (‘when’) component of behavior. PFC might underlie explicit timing through fronto-striatal network (Wiener et al., 2010; Coull et al., 2011; Ohmae et al., 2017) and implicit timing through an interaction with the left inferior parietal cortex (Coull et al., 2016). Human and animal studies show neural signatures of temporal prediction: slow ramping activity that builds while waiting for the expected events and low-frequency rhythms that align behavior with upcoming moments (Xu et al., 2014; Parker et al., 2017; Weber et al., 2024). In humans, EEG recordings identified specific signals called contingent negative variation (CNV) during the time interval (i.e. the foreperiod), which underlie predictive computation (Coull et al., 2011).

The cerebellum, best known for fine-tuning movements, also contributes when behavior needs precise, non-rhythmic timing, and it interacts with PFC through thalamus. Indeed, recent studies showed that cerebello-thalamo-prefrontal loops endow explicit timing in interval-timing discrimination tasks at both sub-second and supra-second timescales (Ohmae et al., 2017; Parker et al., 2017; Breska and Ivry, 2018). In rodents ramping activity in neuronal firing rates were observed both in the cerebellar nuclei and in the PFC (Chabrol et al., 2019; Ren et al., 2025). The cerebellar cortex also stores sub-second timing information in a classical implicit timing task, the eyeblink conditioning, in which primates and rodent learn to close the eyelid following a conditioning stimulus to anticipate an aversive air puff in the cornea (Kim and Thompson, 1997). While learning is essentially cerebellar when the conditioned stimulus and the air puff overlap in time, it is stored in a widely distributed network including the cerebellum, the hippocampus, the amygdala and the PFC when they are separated in time by a delay (Skelton, 1988; Siegel et al., 2015; Weiss and Disterhoft, 2015; Li et al., 2022). Altogether, these studies show that – implicit and explicit - time processing is widely distributed, and thus suggest that information about time might represent a task specific byproduct of network computation.

To test this hypothesis, we designed a simple behavioral task in which two visual cues preceded reward delivery. During training and the first part of the experiments, the interval between the second cue and the reward varied; critically, successful performance did not require temporal discrimination (i.e., timing was implicit), as rewards were non-contingent and failure to respond was not penalized. This design allowed us to examine how time is encoded even when it is not required for task completion. In the final session, the delay was fixed, and we assessed whether mice were sensitive to a slight adjustment in time. We identified anticipatory signatures at both the behavioral and neurophysiological levels, including in mPFC local field potentials and single-unit firing rates. Time was encoded incidentally as variable versus fixed delays elicited distinct neurophysiological correlates. Moreover, optogenetic stimulation of cerebellar Purkinje cells modulated this anticipatory behavior. Together, these findings suggest that time processing in this paradigm is an emergent property of distributed neuronal network dynamics.

## Results

We investigated how the medial prefrontal cortex (mPFC) encodes temporal information during a cued water-delivery task. Water-restricted mice (see Methods) were trained for about 11 days to lick for a water drop following two visual cues. The cues were separated by a fixed 1-s interval, whereas the cue–reward delay was first variable: on each trial the water drop appeared at the spout after either 400 or 900 ms, selected at random (Figure 1A). Mice were head-fixed and could run freely on a non-motorized wheel. Reward delivery was non-contingent (i.e., delivered on every trial), but the drop was withdrawn 150 ms after appearance, encouraging anticipatory preparation to obtain it. Licking was permitted at all times, with no penalties. On the final training day, a single session employed a fixed 500-ms delay to assess sensitivity to a modest time adjustment that did not alter task contingencies. Animals were implanted with multi-unit electrodes in mPFC and an optical fiber positioned above the cerebellar cortex. (Crus I; **Figure 1B; see Methods).**

**Figure 1.**
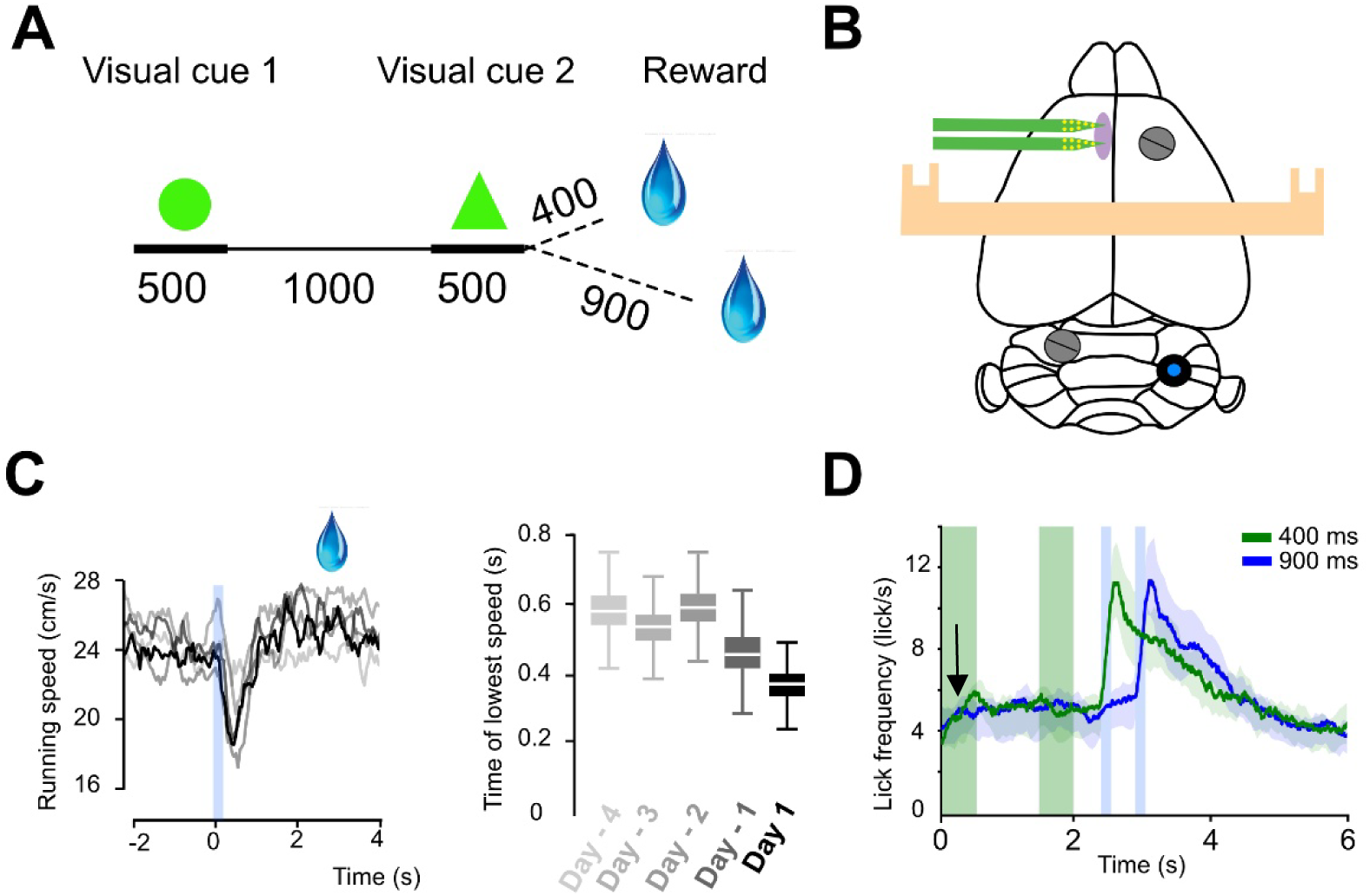
Task description. **A,** experimental design of the cued water delivery task. Two visual cues (0.5 s each, separated by a 1 s interval) precede a water drop delivered after a variable delay (0.4 or 0.9 s). **B**, schematic of the head-fixed setup showing electrode position (green bands), reference and ground screw, optical fiber above the cerebellar cortex (blue circle) and head bar (beige bar). **C**, averaged running speed across training sessions and first day of test (N=10). **D**, averaged licking behavior across trials (N=10). Lick rate increased during presentation of the first cue (black arrow) and peaked sharply at reward delivery.

### Behavioral correlates of reward anticipation during variable delays

We trained mice (N=10) and first quantified running and licking behavior. Running speed was monitored across all sessions and averaged across animals (Methods). After several days of training, mice began to decelerate at trial onset (mean change: –1.27 cm/s; **Figure 1C**) and then showed a pronounced speed reduction at the expected reward time. We used the latency to reach minimal running speed and the lick frequency around reward delivery as proxies for learning. With training, the latency to minimal running speed decreased significantly, reaching a minimum of 366 ± 50 ms, and then returned to baseline following reward delivery (**Figure 1C**). In contrast, licking increased during the first cue, plateaued, and then rose sharply at reward delivery, peaking at 13.3 ± 5.2 licks/s—a 3.8 ± 2.1-fold increase over baseline (**Figure 1D**). Together, these observations indicate that mice not only learned to collect the water drop when it became available, but also adjusted both locomotion and licking from the very start of the trial—consistent with anticipatory behavior during the delay period.

### LFP correlates with anticipation of reward in mPFC during variable delays

Mice were implanted with a 16-channel silicon probe in the left prelimbic (PrL) region of the mPFC, and recordings targeted layer V (AP: +1.9 mm, ML: +0.3 mm, DV: −1.6 mm**; Supplementary Figure 1**), a locus implicated in temporal processing (Parker et al., 2017). L7-ChR2 mice (N = 10)(Chaumont et al., 2013) were implanted before training; two recording sessions were acquired at the end of the training—one with variable cue–reward delays and one with a fixed delay (see **Methods**).

We first analyzed local field potentials (LFPs) during the variable-delay session. For each trial, raw signals were averaged across electrode sites on a shank (**Figure 2A**). Low-pass filtering revealed transient deflections time-locked to the cues and a larger deflection at reward delivery, indicating task-related engagement of prefrontal networks. Time–frequency analysis in individual mice (**Figure 2A**) showed oscillations across multiple frequency bands during the task (e.g., delta and theta; **Figure 2B**). Accordingly, we applied Morlet wavelets on each trial to extract amplitude and phase across frequencies; amplitudes were then band-averaged across trials within a session and subsequently averaged across mice (**Figure 2B**). Amplitudes increased significantly at reward time for both delays. In addition, delta, theta, and beta amplitudes exhibited a gradual rise from trial onset (**Figure 2C**), consistent with mPFC processing of task-relevant information during the delay.

**Figure 2.**
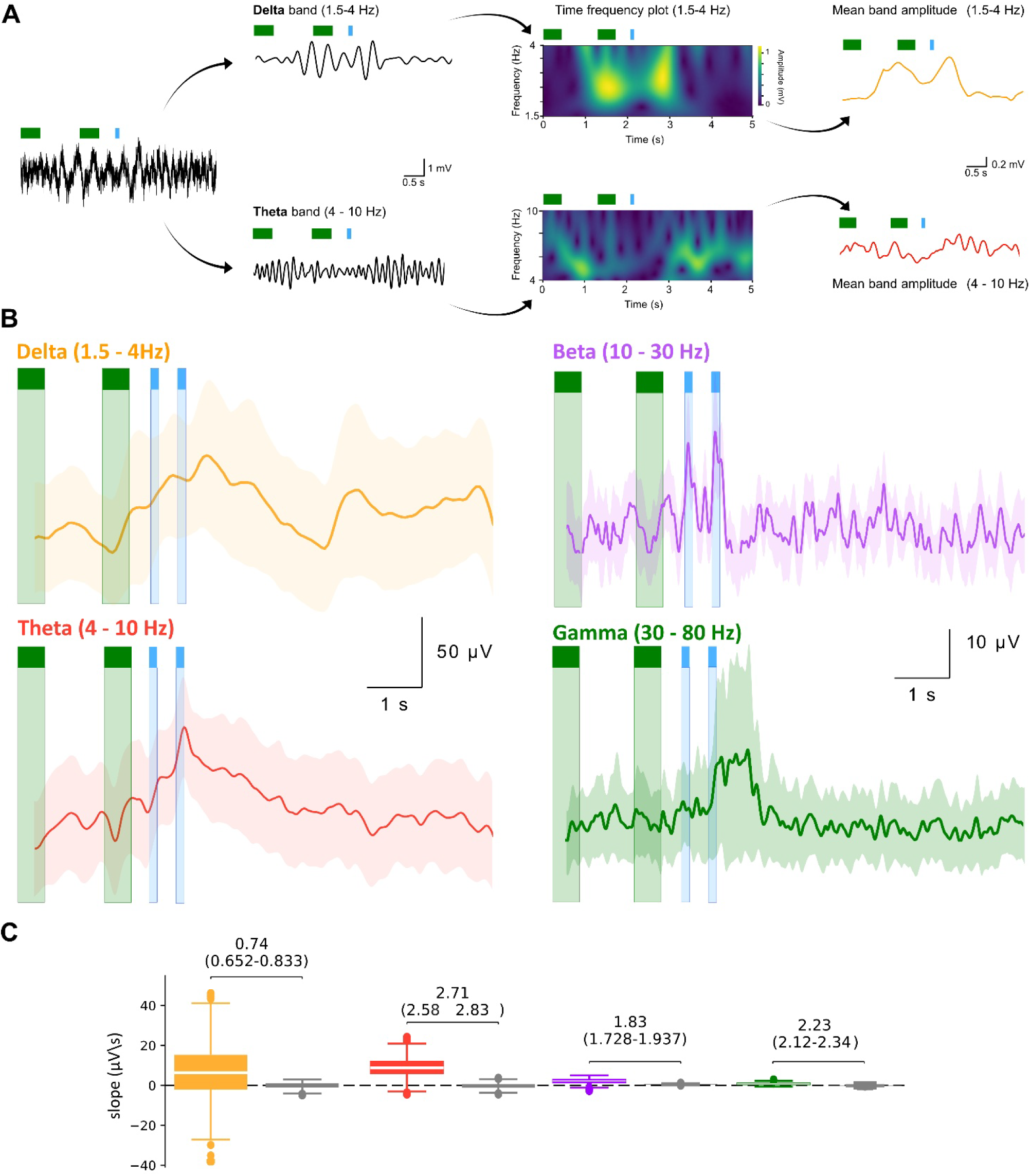
Task-related oscillatory dynamics in mPFC. **A,** example of preprocessing and analysis applied to raw LFP signals. The signal was first band-pass filtered in the frequency range of interest, then transformed into a time–frequency representation using Morlet wavelet analysis (see Methods). Amplitudes were then band-averaged across trials within a session and subsequently averaged across mice. **B,** Time course of averaged band amplitudes across mice (N=9). **C,** bootstrapped distributions of the slope of the averaged amplitudes computed from the anticipatory period for each band, compared with a permutation-based null distribution generated by time-resampling within trials (N=9). Reported values correspond to Cohen’s d effect size, quantifying the strength of the anticipatory increase in LFP amplitude relative to this null distribution.

To assess phase alignment, we computed inter-trial phase consistency (ITPC) as a measure of phase concentration (**Methods**), where ITPC = 1 indicates identical phase across trials at a given time point. Delta, theta, and beta phases showed pronounced synchronization at reward delivery (**Figure 3**), indicating a reward-locked phase reset of mPFC oscillatory activity.

**Figure 3.**
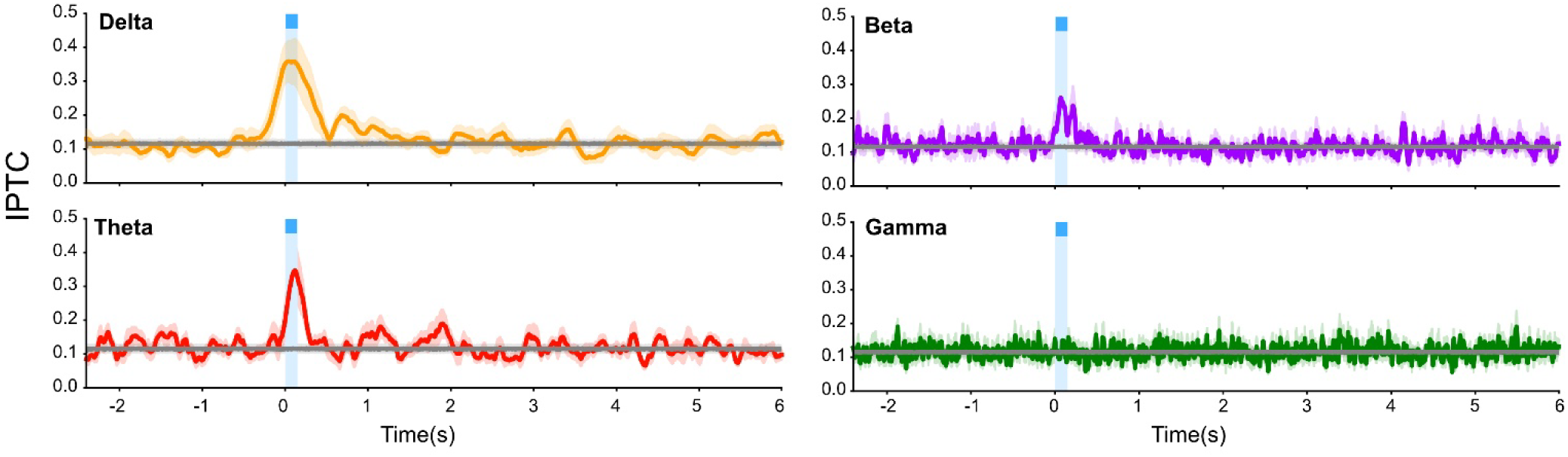
Inter-trial phase consistency (ITPC) of oscillatory activity in the mPFC. Mean ± s.e.m. of inter-trial phase consistency (ITPC) for delta, theta, beta a,d gamma bands recorded in mPFC. Phase alignment peaked at reward delivery, suggesting a trial-by-trial phase reset and synchronization of network oscillations by reward-related input. In grey, mean ± s.e.m. IPTC of shuffled data.

### Phase-locked pyramidal neurons in mPFC are more active and have specific dynamics

Spike sorting (**Figure 4A**, **Supplementary figure 2**, **Methods**) allowed us to discriminate individual units including a majority of putative pyramidal cells (n = 106 pyramidal neurons vs 7 interneurons, N = 8 mice). In the following, only pyramidal cells were analyzed. We assessed whether individual units are correlated with brain oscillations identified in the LFP. Phase locking was computed by registering the spikes from a unit to the position in the phase for each frequency band, and the distribution was tested for uniformity (Rayleigh test, **Figure 4B**). We showed that almost all neurons (84 %, n = 88) were phase-locked to the gamma band, suggesting that layer V pyramidal cells are engaged in local network processing. Moreover, neurons phase-locked to the gamma band have a several fold higher firing rates compared to non-phase-locked neurons (mean firing rate phase locked neurons = 4.6±3.9 Hz vs 0.7±1.1 Hz, **Figure 4B**). In the other frequency bands, we identified 34%, 25% and 29% neurons phase locked to beta, theta and delta band, respectively (**Figure 4B**). While in the beta and theta frequency bands phase-locked neurons have also a significant higher firing rate (Theta phase locked = 5.3±3.3Hz vs 3.5±3.9Hz; Beta phase locked = 5.9±4.4 Hz vs 3.1±3.3 Hz), it was not the case in the delta band. Oscillations in the delta band have been associated with long-range communication with distant brain areas such as the cerebellum (Parker et al., 2017; Breska and Ivry, 2020). Therefore, these results suggest that in variable delay tasks, long range communication might be preferentially mediated by beta and theta band. While averaging all pyramidal neuron firing rates yielded a weak correlation to the task (**Supplementary Figure 3**), individual averaged firing rates of individual unit showed modulation locked to the different features of the task with units ramping up or down during the delay, and peaking at the reward time (**Figure 4A**).

**Figure 4.**
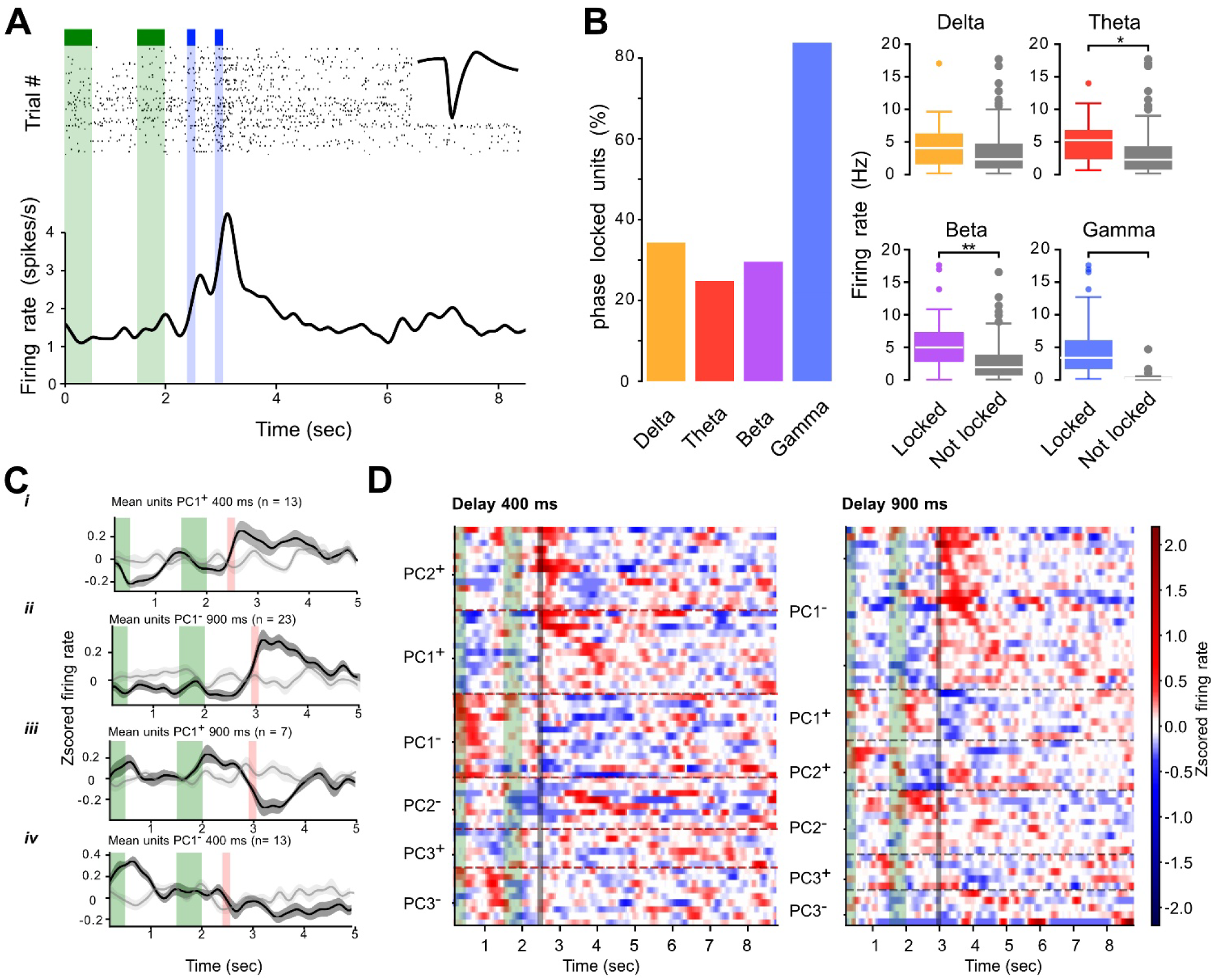
Temporal dynamics of single-unit activity in mPFC. **A**, example of raster plot of a single unit and its averaged waveform (inset). *Bottom*, mean firing rate of the unit during the session (see **Methods**). **B**, *left*, proportion of pyramidal neurons phase-locked to each frequency band and their mean firing rates, 34%, 25%, 29% and 84% of neuron are phased lock to delta (locked n=36, not locked n=73), theta (locked n=32, not locked n=83), beta (locked n=31, not locked n=78) and gamma (locked n=88, not locked n=21) bands, respectively. *Right*, phase-locked neurons showed significantly higher firing than non-locked units, in theta (p-value = 0.03, t-test), beta (p-value = 0.002 t-test) and gamma (p-value = 1,3.10^-11^, t-test) range but not in delta range. **C**, mean normalized firing rate of clusters of units identified by principal component analysis (PCA). **D**, heat map showing normalized firing rate of individual neurons (each row = one unit), sorted by principal components (n=56 for 900ms and n=62 for 400ms). Horizontal dotted lines indicate cluster boundaries. Green and black vertical bands identify cue and water reward timing, respectively.

We also observed units for which the second cue was correlated with clear alteration of the mean firing rate (**Supplementary figure 4**). Using principal component analysis (PCA), we therefore set out to identify latent variables based on firing rate behaviors in the pyramidal cell population (**Figure 4C-D; methods**). This analysis was performed on trials where the reward was delivered at 400-ms and at 900-ms separately. While 12–13 principal components were required to explain 95% of the variance, latent variables were considered relevant only when the firing rates of at least five units correlated with a principal component (p-values < 0.05; see Methods), yielding three main principal components (41.8% and 40.6% explained variance for the 400-ms and 900-ms conditions, respectively, corresponding to 56/109 units and 62/109 units). We then identified and plotted the average firing rates of neurons that were correlated or anti-correlated (−0.5 < r < 0.5) with these main principal components. Three major categories of firing-rate time courses were observed: (1) those peaking high or low at reward time, (2) those ramping up or down during the delay, and (3) those peaking at the moment of the second cue. In all conditions, the second cue and the reward time appeared as key moments during the behavior. Altogether, we show that individual neurons are embedded in local (gamma band) and external (beta, theta band) oscillatory activity, and that both oscillations and units correlate with the behavioral task. Furthermore, a low number of orthogonal latent variables encode the task in the mPFC. Taken together, these results suggest that, although the precise timing of the reward is not fully predictable, neuronal networks in the mPFC learn to anticipate reward delivery.

### A single fixed delay session leads to fast adaptation of behavior

We then asked whether making the delay fixed—and therefore more predictable—would alter behavior and neurophysiological measures. We used a single fixed delay (500 ms) for a single day session of the experiment, the last session. At the end of this session (60 trials), we observed that behavioral anticipation was significantly affected with a steeper increase in licking behavior during the delay (0.85 ± 1.08 lick.s^-1^ vs -0.03 ± 0.43 lick.s^-1^ for fixed and variable conditions, respectively; p-value = 0.023 paired t-test; N=10; **Figure 5B**). Neither the timecourse of the amplitude of oscillations nor the overall mean firing rate of recorded units (4.4 ± 4.7Hz vs 4.1 ± 4.1Hz for fixed and variable conditions respectively, n = 109 and n = 115) were significantly affected by the fixed delay (**Supplementary Figure 5**). However, while the firing rates of phase locked units to the delta band were indistinguishable from non-phase locked units in variable delays (4.7 ± 4.1Hz n = 38 vs 3.8 ± 4.1Hz n = 71 for phased locked and not phased locked respectively, **Figure 4B**), firing rates of delta band phase-locked units were significantly increased during the single fixed delay session (p-value = 0.032, t-test; 7.0 ± 5.2Hz n=16 vs 3.9 ± 4.4Hz n=97 for phased locked and not phased locked, respectively; **Figure 5C**), albeit their number decreased (from 34.3% in the variable condition to 14.3% in the fixed condition). Spiking activity under fixed-delay conditions showed similar categories of firing rate time courses as in the variable-delay condition: (1) peaking high or low at reward time, (2) ramping activity during the delay, and (3) cue-aligned peaks (**Figure 5D**). A PCA performed on averaged firing rates of individual units as above showed that these categories were present in comparable proportions, indicating that the same basic firing patterns were preserved across conditions. However, while peaks in IPTC were also observed at reward time, phase alignment significantly increased in the beta band oscillation compared to variable delays (**Figure 5E**). Altogether, since delta and beta oscillations are associated to long range communication pathway (e.g. cerebello-cortical) and to the assessment of novelty both in mice and human (Breska and Ivry, 2020; França et al., 2021), these results suggest that mPFC units may receive stronger long range inputs in the fixed delay protocol. Overall, we postulate that increasing delay predictability during the anticipation phase (randomly chosen 400–900 ms vs a fixed 500 ms) within a single session in this implicit task can shape both behavior and mPFC network synchronization.

**Figure 5.**
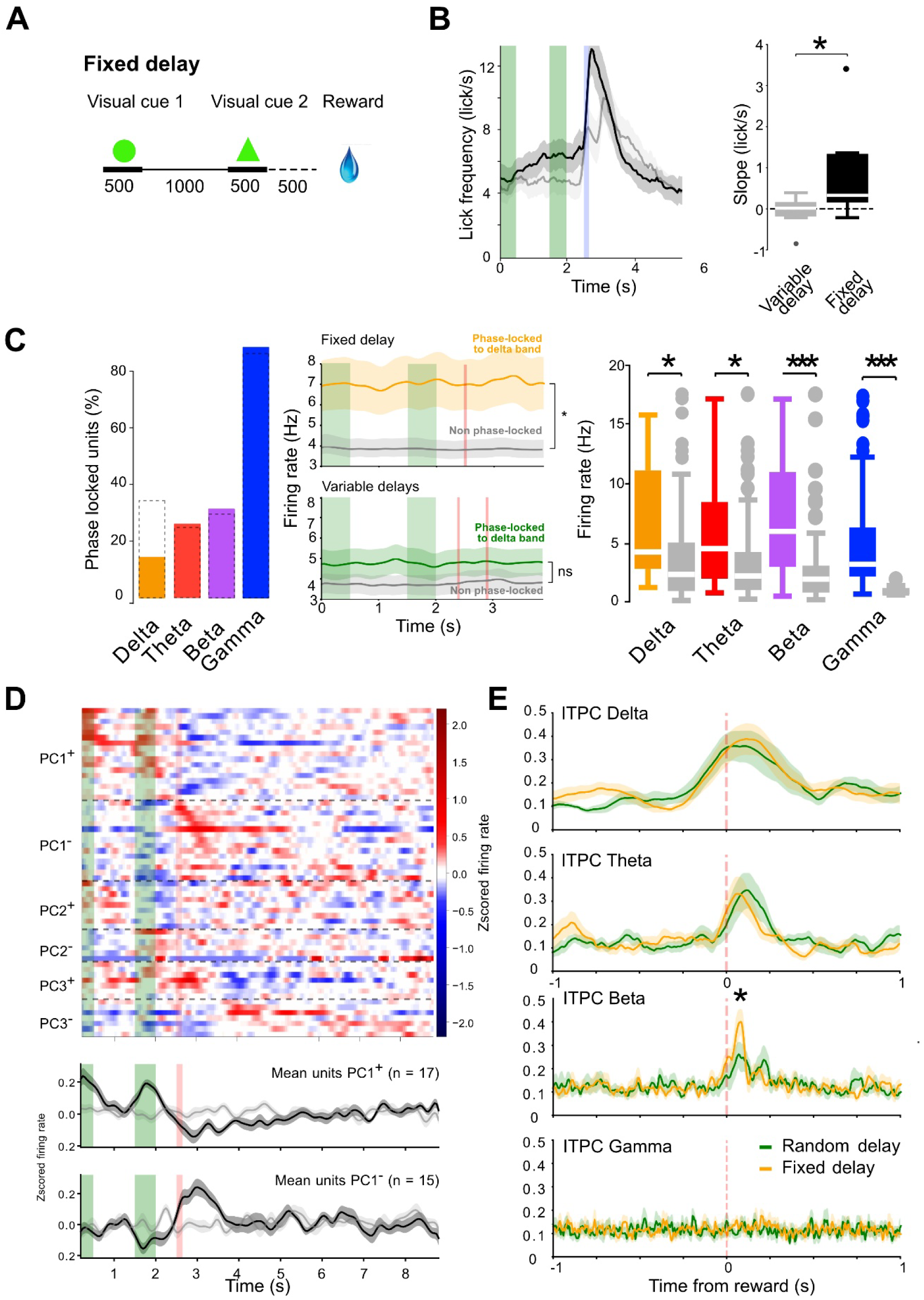
Adaptation to a fixed delay. **A**, experimental design of the fixed-delay session (0.5 s). **B**, mean lick frequency and boxplots of the slopes of the anticipatory phase (first cue to reward time), N = 10. **C**, *left*, histogram of phase-locked neurons to delta, theta, beta and gamma bands. *Middle*, mean firing rates of all phase locked units to delta band in variable and fixed delays compared to non phase-locked units (p-value = 0.032, t-test; 7.0 ± 5.2 Hz n=16 vs 3.9 ± 4.4 Hz n=97 for phased locked and not phased locked, respectively). *Right*, boxplots of phase-locked units per band compared to non-phased-locked (delta locked units n=16, not locked units n=97, p-value = 0.044 t-test; theta units locked n=29, units not locked n=84, p-value = 0.014 t-test; beta units locked n=35, units not locked=78, p-value =4.5×10^-5^; gamma units locked n=99, units not locked n=14, p-value = 8.0×10^-15^). **D**, *upper*, heat-map of the units sorted after Pearson correlation with principal components of the PCA (PC1-3, n=61). *Lower*, averaged z-scored ± s.e.m. firing rates (black) of units positively and negatively correlated to the first principal component compared to shuffled spike train (grey). For PC1^+^, firing rates were significantly different between 0 and 1.5 sec (F=8.8 p-value = 0.009, RM ANOVA), 1.5 and 2.5 sec (F=9.7 p-value = 0.007, RM ANOVA), and 2.5 and 4 sec (F=25.3, p-value = 0.0001, RM ANOVA). For firing rates were significantly different between 1.5 and 2.5 sec (F=7.8 p-value = 0.014, RM ANOVA), 2.5 and 4 sec (F=18.2, p-value = 0.0008, RM ANOVA). **E**, comparison of averaged ITPC between fixed and variable delays (N=9; *p-value = 0.035, Kolmogorov–Smirnov test).

### Cerebellar stimulation influence mPFC cortical dynamics and behavior

We examined long-range communication between the cerebellum and the mPFC using optogenetic activation of cerebellar Purkinje cells (PCs). To assess functional connectivity between the two regions, we first stimulated PCs in a cohort of anesthetized L7–Channelrhodopsin-2 mice (L7-ChR2) mice (Özcan et al., 2020). Extracellular event-related potential recordings were obtained from the left Prelimbic (PrL) while brief blue-light pulses were delivered to defined sites on the right cerebellar surface (**Supplementary Figure 6**). Illumination of right crus I produced a significant deflection in PrL local field potential (LFP) amplitude; accordingly, subsequent experiments in awake animals targeted this region, consistent with prior work (Parker et al., 2017; Kelly et al., 2020).

For awake recordings, mice were implanted with a multiunit recording electrode in the left mPFC and an optic fiber positioned above right crus I before training. Across two recording days, a single series of photostimulation (115 Hz; 30 trials) was delivered from the end of the second cue to reward delivery at an intensity of 16 mW/mm²). We previously showed that photostimulation elicits a significant increase in PC firing (Chaumont et al., 2013) yielding both a reduction and an entrainment of cerebellar nuclei projection neurons (Özcan et al., 2020).

Photostimulation of PC had different effects in the variable and fixed delay conditions. In the variable 900 ms delay condition, Purkinje cell activation accelerated locomotor responses: locomotion slowed prematurely on these trials, and the time course of this effect closely matched the behavior normally observed in the 400 ms trials (**Figure 6A**). Mean firing rates also increased selectively in the 900-ms trials and reached their maximum at the earliest possible reward time (**Figure 6B**). In parallel, stimulation induced an early rise in theta ITPC in the same 900-ms trials and asignificant effect in the gamma band (**Figure 6C).** In the fixed-delay condition, licking behavior was affected, with a significant reduction in lick rate at reward delivery; we also observed decreases in delta and beta ITPC (**Figure 6C**). Taken together, the stimulation influenced both behavior and mPFC network dynamics, but the contrasting outcomes between the variable and fixed delay conditions indicate that each timing regime may engage distinct mPFC networks or network states.

**Figure 6.**
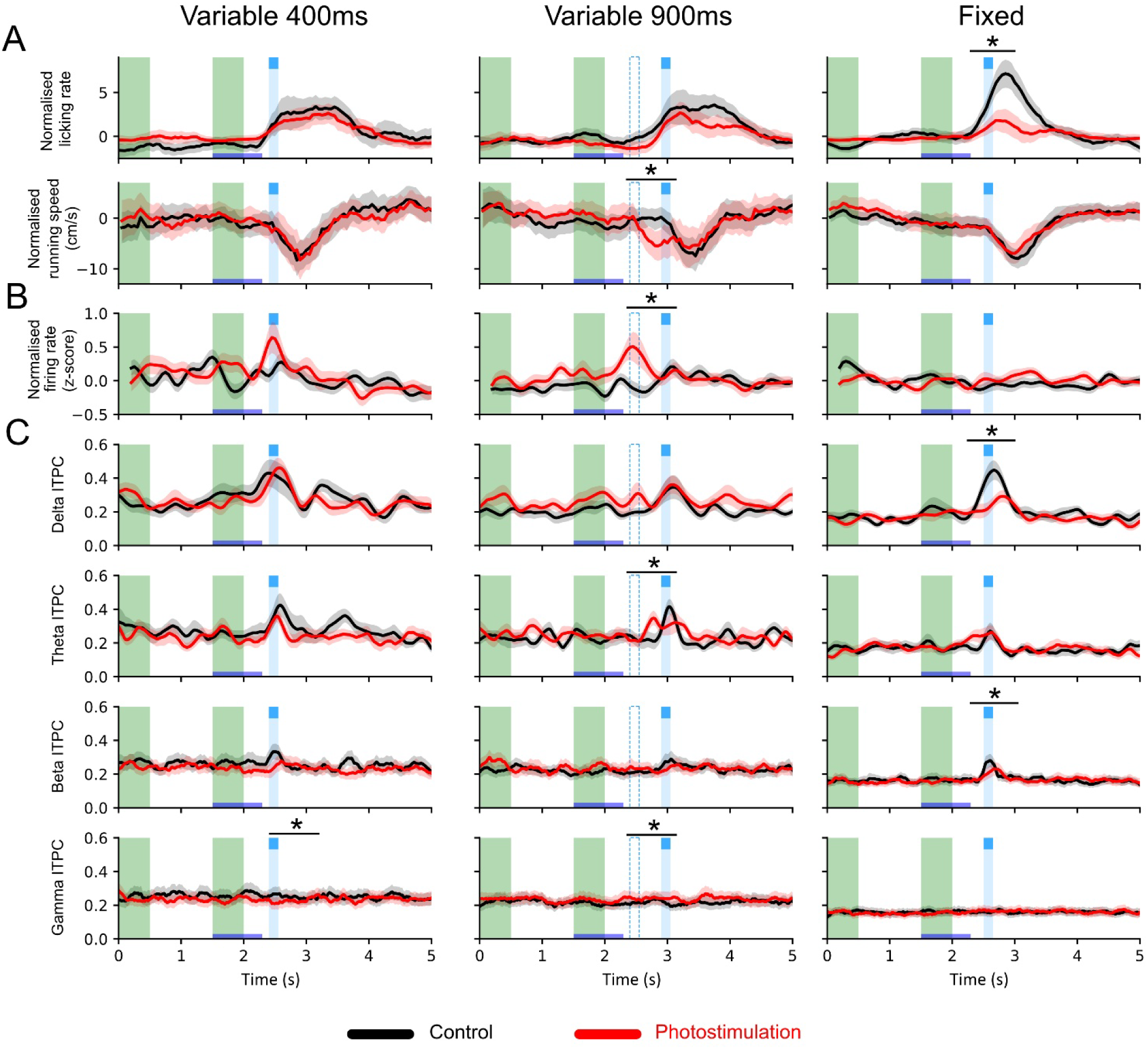
Purkinje cell stimulation affects behavior and mPFC networks in a delay-dependent manner. **A, *upper,*** Licking behavior during optogenetic stimulation of Purkinje cell in Crus I (115 Hz, 16 mW) in variable delay and fixed sessions. *Lower*, running speed during photostimulation. **B**, averaged neuronal firing rate in the mPFC during photostimulation. **C**, Inter-trial phase consistency (ITPC) in delta, theta, beta and gamma bands. Two-way repeated-measures ANOVA were applied to each metric from the onset of the first possible reward at 2.4 s to the end of the last possible reward at 3.05 s. *indicate significance levels; panel A, running speed at 900 ms F = 7.7, p- value = 0.022, n=10, licking firing rate fixed delay F = 9.0, p-value: 0.015, n=10; panel B, interaction F = 5.1, p- value = 0.011, n=110; panel C, delta IPTC fixed delay F= 19.2, p-value = 0.002, interaction F = 9.0, p- value = 0.004, n=9, Beta IPTC fixed delay interaction F = 3.0, p- value = 0.018, n=9; Theta IPTC 900 ms, interaction F = 3.3, p- value = 0.027, n=9; gamma IPTC, F = 15.7, p-value = 0.004, n=9 for 400 ms, and F = 5.4, p-value = 0.047, n=9 for 900ms.

To quantify the effects of photostimulation at the level of individual units, we used a classifier-based approach to test whether trials with cerebellar stimulation could be discriminated from non-stimulation trials using only neuronal firing rate (**Figure 7A**). For each time bin of the task (50 ms bin width), a classifier was trained to discriminate between stimulation and non-stimulation conditions using the firing rate of each recorded unit. Classifier performance was quantified by the area under the receiver operating characteristic curve (auROC), where a value of 1 indicates perfect discrimination and 0.5 corresponds to chance level. The mean auROC across units was significantly above chance (variable 400 ms: 0.546 ± 0.08 SD, p-value = 2.4×10^-11^; variable 900 ms: 0.551 ± 0.083 SD, p-value = 2.9×10^-7^; fixed: 0.575 ± 0.102 SD, p-value = 4.8×10^-9^, paired t-test, n=113), indicating that trial identity could be reliably decoded from neuronal activity at almost all time bins (**Figure 7A, B**). In many units, the best decoding scores occurred near the first possible reward time (2.4 s), where auROC values reached their maximum (**Figure 7A, B**). A hierarchical clustering of individual units based on auROC timecourses revealed that the cluster of units having the best accuracy do not belong to any subtypes of firing rate profiles identified by PCA (see **Figure 4C, D**) but were distributed across all the profiles. Together, these results indicate that cerebellar photostimulation influenced mPFC at the single-trial level, with discriminative activity concentrated around the earliest possible reward time. However, significant decoding accuracy was observed beyond this period, suggesting that cerebellar stimulation induced both transient and persistent modulations of cortical activity.

**Figure 7.**
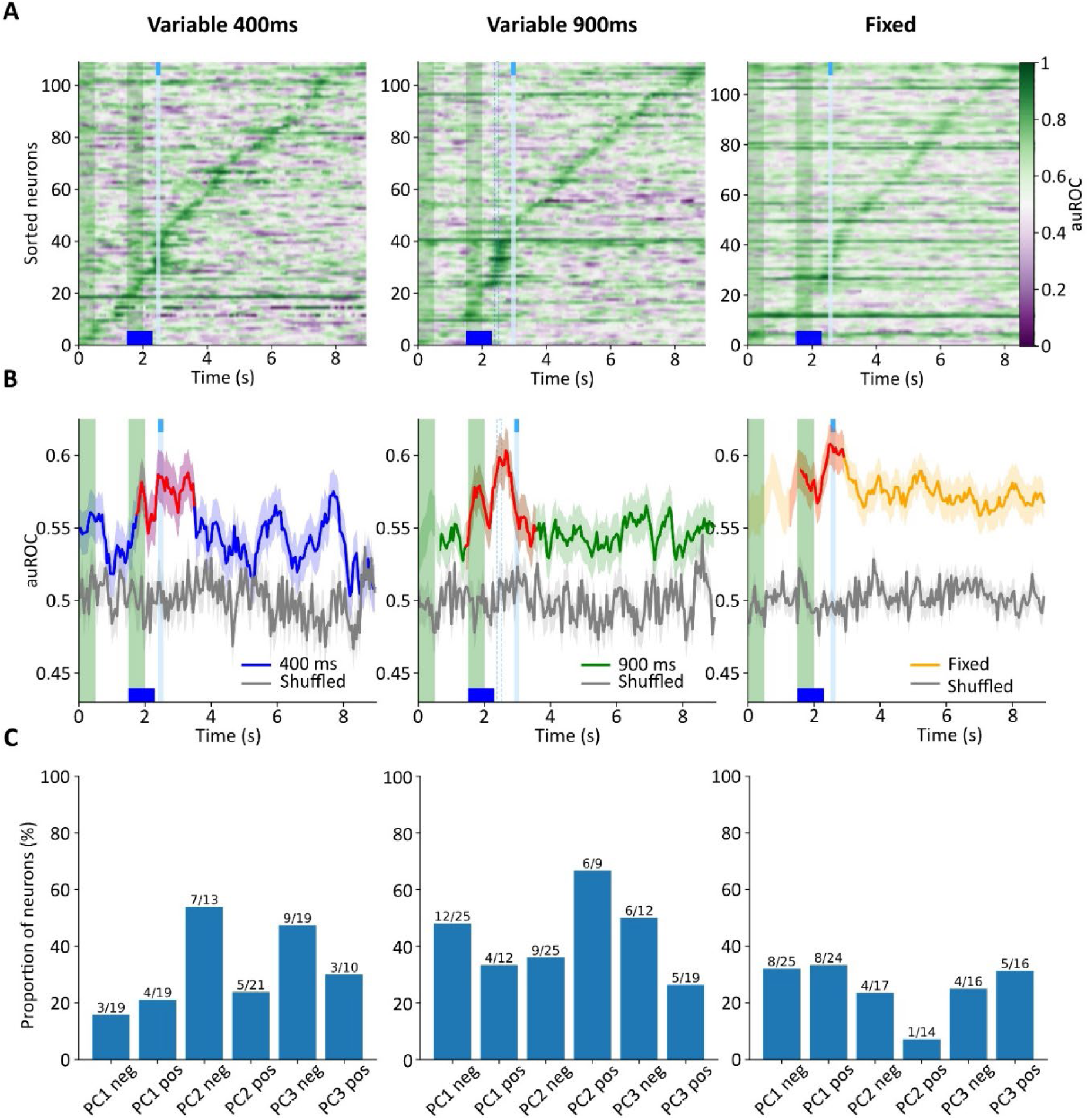
Classifier-based decoding performance between stimulation and non-stimulation trials. **A**. Time-resolved decoding accuracy computed as the area under the receiver operating characteristic curve (auROC) for each time bin and neuron. Higher values indicate higher discriminability between trials with and without cerebellar Purkinje cell stimulation, with 0.5 being chance level. **B**. Averaged timecourse of decoding accuracy from units shown in A compared to shuffled data (in grey, see methods). In red, time bins with the highest accuracy identified by PCA (see **supplementary Figure 7**) **C.** Distribution of Units identified Neurons whose maximal auROC values fell within the extracted epoch were examined to determine whether the effect reflected specific population features or was broadly distributed across PC clusters (see Figure 4). The results indicate that the effect was distributed across clusters. Green bands correspond to cue presentations and the blue line marks the time of reward delivery.

## Discussion

### Time encoding is an emergent property of the mPFC network

We here demonstrated that mice can learn to anticipate a cued reward when the delay between the cue and the reward is variable. In this task, there was no punishment, and the reward was delivered during all trials, but only during 150 ms. Therefore, although the interval between the cue and reward provides little temporal information, mice must be ready to lick to catch the water drop. For this reason, we considered that timing was implicit in this task. Most previous works focused on an explicit version of interval timing in which animals have to discriminate two intervals to get rewarded (i.e. temporal bisection task; (Tallot and Doyère, 2020). While we acknowledge that after training the task developed in this study may be considered as explicit once mice become experts, perception of time was still implicit as it had no relevance for the outcome of the trials (no success nor failures). Anticipation was observed at the behavioral level as mice slowed down at reward while intensifying licking as observed in preparatory activity (Chabrol et al., 2019). Furthermore, neuronal network in mPFC encoded the task in LFP and in specific firing rates dynamics in layer V pyramidal cells. Our recording device using 16 channels silicon probes did not allow dense mappings in individual animals, we therefore did not attempt to correlate latent variables to specific features of the behavior in individual animals. However, by recording the output layer of the mPFC, we could extract several general neuronal timecourses in pyramidal cells. Too few interneurons were recorded and were consequently discarded from our analysis. Overall, while about 6 main profiles have been identified, two main features appeared salient and robust: (1) individual units ramped up or down before the reward and peaked at the time of the reward, and (2) unit firing rates are deeply influenced by the second cue. These two features are in agreement with previous work using longer and explicit time intervals suggesting that time encoding is a fundamental property of the mPFC, which may arise as an emergent network processing (Oshio et al., 2006; Kim et al., 2013; Xu et al., 2014; Bakhurin et al., 2017; Parker et al., 2017). Furthermore, time encoding is quickly adjustable and flexible as a single session at a different and fixed delay influenced both behavioral and neuronal features. Indeed, instantaneous temporal adjustments in temporal updates during behavioral tasks have been already widely documented when intervals were previously experienced by the mice (Li and Dudman, 2013; Aggadi et al., 2025). Since the fixed delay in our experiments is close to one of the variable delay (500 vs 400 ms), we postulate that our results suggest that mice associate the fixed delay to the shortest of the variable delay they were trained with.

### Modifying predictability quickly affects time encoding

Importantly, while the absolute timing of reward delivery was similar across conditions, the level of predictability differed. Our data indicate that mice were sensitive to this change and appeared to adjust their behavioral strategy, as suggested by the steeper anticipatory licking slope under the fixed-delay condition. In the variable condition, one plausible interpretation is that mice focused on discriminating between the two possible reward times and continuously updating whether the reward had already occurred, whereas in the fixed condition they may have been able to encode the expected reward time more directly and adopt a more strongly anticipatory strategy as observed in humans (Coull et al., 2016), which is reflected behaviorally in the steeper licking slope. Optogenetic stimulation of Purkinje cells further emphasized these condition-dependent differencesIn the variable-delay condition, perturbing cerebellar activity did not abolish reward-related responses; instead, it advanced several aspects of task-related dynamics, including an earlier reduction in locomotor speed and increased firing around the earliest possible reward time. This pattern is consistent with interference in ongoing temporal updating and with the cerebellum’s established role in movement initiation (Kunimatsu et al., 2018; Dacre et al., 2021). In the fixed condition, by contrast, the same perturbation was associated with a reduced reaction to reward, evidenced by a lower licking rate after reward delivery and a decrease in delta and beta band ITPC, compatible with disrupted anticipation. Notably, beta bands ITPC was significantly affected in the fixed condition, in line with our observation that fixed delay specifically modified the coupling of neuronalactivity to betaoscillations and with an identified role of beta oscillations in time processing (Kulashekhar et al., 2016; Stoll et al., 2016). This also aligns with previous evidence that delta rhythms are critical for interval timing and provide a pathway through which the cerebellum can modulate prefrontal activity (Arnal et al., 2015; Parker et al., 2017; Herbst and Obleser, 2019). Together, these results indicate that mice are sensitive to implicit changes in temporal predictability, and that this sensitivity is reflected in distinct states of the cerebello-prefrontal network (Siegel et al., 2012). In addition, the decoder showed that the different effects of photostimulation in the variable and fixed conditions were not due to a failure to alter local mPFC activity, as auROC values remained above chance in the fixed condition. Instead, this suggests that the effect was timing-dependent and selectively modulated event-related responses linked to processing the expected reward time. An alternative explanation is that the observed effects reflect learning-dependent dynamics. Since the fixed delay occurred in a single session of 90 trials after several days of training with variable delays, mPFC dynamics adapted during this session, and led to a modified sensitivity to cerebellar stimulation. Indeed, many studies showed that the cerebello-thalamo-cortical loop is involved during the learning phase of a task (Galea et al., 2011; Pemberton et al., 2024), which could explain the differential effect of the stimulation during the fixed delay condition.

### Low-frequency oscillations mediate cerebello–PFC communication

Our results agree with other studies showing that specific oscillatory bands underlie communication between the cerebellum and the forebrain in the context of time processing. Rodents learn to discriminate interval of time both at the second or sub-scond timescale. In these paradigms, accurate discrimination correlates with an increase in the amplitude of delta band oscillations during the interval. In a pharmacological model of schizophrenia, explicit interval timing discrimination is impaired and the delta band vanishes, but during cerebellar optogenetic stimulation the accuracy of the discrimination recovers in coincidence with an increase in delta band oscillations. Interestingly, similar delta band failure during interval timing discrimination was observed in individuals with schizophrenia (Parker et al., 2017). Furthermore, an alteration of connectivity in the dorso-lateral prefrontal cortex - cerebellar network is directly linked to negative symptom severity in schizophrenia (Brady et al., 2019). In these patients, rTMS stimulation to the cerebellar midline in humans alleviate negative symptoms via an adjustment of the cerebello-prefrontal communication indicating that the cerebellum can entrain prefrontal networks. In Breska and Ivry (Breska and Ivry, 2018, 2020, 2021) studies on interval timing in cerebellar degeneration disease, the authors used the inter-trial phase concentration parameter, a metric similar to the ITPC used in our study to assess how frequency bands synchronize during the task. They demonstrated that the scalp topography of delta band inter-trial phase concentration was associated with target anticipation, with the control group exhibiting stronger inter-trial phase concentration than the cerebellar degeneration group in the interval task. This suggests that the delta band activity is involved in forming temporal predictions based on single intervals, which is impaired in individuals with cerebellar dysfunction. The beta-band plays a different role in interval timing. While the delta-band activity is associated with target anticipation in the interval task, the beta-band activity is suppressed at the time of an expected event that requires a motor response, and this feature is impaired in individuals with cerebellar degeneration in interval-based predictions, but not in rhythm-based predictions (Breska and Ivry, 2020).

### Comparison with eyeblink conditioning

In most previous studies interval timing was explicit and supra-seconds, with animals being rewarded for successes (Xu et al., 2014; Parker et al., 2017). Here, we demonstrate that interval timing is processed incidentally as a non-conditional by product of task learning by the brain. Our task has some similarities with the eyeblink conditioning (EC) task (McCormick and Thompson, 1984; Kim and Thompson, 1997; Yeo and Hesslow, 1998; Johansson et al., 2016). This is a classical implicit timing task in which primates and rodents learn to close the eyelid following a conditioning stimulus (CS) to anticipate an aversive puff in the cornea (US for unconditioned stimulus) after repeated pairing of the CS and US. When the interval between CS onset and US overlap and are in the sub-second range, the cerebellar cortex stores the temporal information (i.e. Delay Eyelid Conditioning). Conversely, when the CS and the US do not overlap and are separated by a delay exceeding one second (i.e. Trace Eyelid Conditioning), the temporal information behaviour is controlled and stored in a distributed network encompassing the hippocampus and the cerebello-thalamo-prefrontal pathway. Perturbation in the cerebellar cortex, the thalamus or in the mPFC area impair these implicit timing tasks (Heiney et al., 2014; Siegel et al., 2015). While the task developed in our study may appear similar to EC, an important difference is that the air puff used during EC is an aversive stimulus, which is a highly salient stimulus compare to the visual cues in our paradigm. We yet observed that the anticipatory behavior on our experiment was learnt in a few days, a similar timecourse than EC. Therefore, interval timing processing is quickly learnt without requirement of a punishment.

### Cerebellar Crus I area is involved in time processing

In our study, we choose to stimulate PCs in crus I, the hemispheric part of cerebellar lobule VII, because several studies identified bidirectional communication between crus I and PrL in the mPFC (Badura et al., 2018; Kelly et al., 2020). Indeed, we demonstrate that optogenetic stimulation in anesthetized animals elicited the response in the LFP in PrL. The cerebellum does not project directly to the cortex but via a disynaptic pathway through multiple pathways that relay either in the ventral tegmental area (VTA; (Carta et al., 2019)) or in the mediodorsal nucleus of the thalamus (MD; (Frontera et al., 2023)) or in the ventromedial thalamus (VM; (Kelly et al., 2020)). Since crus I projects essentially in the lateral cerebellar nucleus, we postulate that the contribution observed in our study is mediated via VM. These pathways are functionally very diverse, the first one operating through dopamine (Rogers et al., 2011), while the others are involving distinct thalamo-cortical subtypes (Clascá, 2022) which target distinct cortical subcircuits (Anastasiades et al., 2021). Whether and how these circuits are engaged in temporal processing is currently unknown; so far, evidence indicates the contribution of the cerebello-VM/VTA-prefrontal cortex pathways in social interactions (Kelly et al., 2020) while the cerebello-MD-prefrontal cortex pathway has been involved in the regulation of emotional state and learning (Frontera et al., 2023). How these circuits are integrated in time-processing remains to be established.

## Methods

Experiments were conducted at the chronobiotron (UAR3415) in accordance with the guidelines for the care and use of Laboratory animals and the French national law (implementing the European Union Directive 2010/63/EU), and they were approved by the regional Ethical committee of Strasbourg (CREMEAS) and authorized by the French Minister of Higher Education, Research and innovation (APAFIS N°36473-2022020711387224 v6).

### Mice

3-4 months old male and female L7-ChR2-eYFP Mice (Chaumont et al., 2013) expressing channelrhodopsin2 in cerebellar PC and bred under the CD1 outbred background were used. Mice were housed 3/5 per cage with littermates of the same sex, with food, water ad libitum and a 12h light/ 12h dark cycle.

### Surgeries

Mice were anesthetized with Isoflurane (Vetflurane, Virbac, 3% for induction and 1.5-2.5% for maintenance) mixed with O2. Mice were placed on the stereotaxic frame (model 68526, RWD Life Science). Body temperature was monitored through a rectal probe connected to a heat-pad (TC-1000, Bioseb Lab) and kept at 37 °C. Before starting the surgery, we implement a multi-modal analgesic pain management to prevent pain and inflammation, with AINS treatment (Metacam (2 mg/kg), injected intraperitoneally) combined with a sub-cutaneous injection of a mix of local local anesthestics, bupivacaine and lidocaïne (2mg/kg each) 2 minutes before skull incision. Craniotomies were performed with a manual driller (OmniDrill35 Micro Drill, World Precision Instruments). A silicon probe (Buzsaki16-CM16LP, Neuronexus) was implanted in layer V of the left prelimbic cortex (PrL; in mm, AP: +1.9, ML: +0.3, DV: -1.6). DV coordinate was calculated from the surface of the cortex. The probe was soaked in DiI (Vybrant DiI cell-labeling solution, Invitrogen, ThermoFisher Scientific) for histological validation. Biocompatible silicone (Kwik-Cast Silicone Sealant, World Precision Instruments) was used to close the craniotomy. Two stainless steel screws (DIN84 M1×2) were implanted as ground and reference to the probe, above left cerebellum, and right frontal area, respectively. A craniotomy was also performed above the cerebellar cortex and a cannula with an optic fiber (zirconia cannula, Prizmatix) was implanted on the surface of the cerebellar cortex (right Crus I, AP: -6, ML: -3). The head-bar (custom made, glass fiber) was sterilized and glued to the skull, between Bregma and Lambda, using Super-Bond C&B (Sun Medical). After a first layer of cement cured, a final layer of dental cement (Paladur, Kulzer) was applied to secure the stability of the implants. 200 µL of sterile NaCl 0.9% was injected intraperitoneal for hydration. After surgery, mice were left to recover in an isolated cage under red light heat-lamp for a few hours until they were fully awake and then placed back in their home cage with littermates. For the 3 days following the surgery, mice had Oral Metacam (5 mg/kg) diluted in the water supply. They were allowed to recover for a full week before the beginning of the water restrictionprotocol.

### Histology

At the end of the experiments, mice were anesthetized with Isofluorane and euthanized with Euthasol (400 mg/kg). Implants (electrode and optic fiber) were removed and positions were marked through the craniotomy. The brain was dissected out and soaked in in paraformaldehyde 4% (PFA 4%) for 24 hours. Brain slices were performed using a vibratome (Leica VT1000, Germany), mounted and observed with epifluorescence.

### Head restrained setup

The behavioral setup is composed of a commercial part (a wheel and a head-restraining apparatus, Imetronic, France) and of a custom-made part (LED control and reward delivery system using routines written in LabView, National Instruments). Wheel speed was monitored using 16 checkpoints along the perimeter of the wheel and logged using the LabView interface. Reward (a drop of water) delivery was controlled by the same routine via a TTL triggering a solenoid valve (Series 3 – Miniature Inert Liquid Valve, Parker). The water was delivered through a spout inserted into a tube connected to a vacuum pump ensuring drop removal after a given duration. Licking is detected when the tongue crosses a light beam.

### Recordings

Chronically Implanted silicone probes with 16 channels (2 shanks, 8 sites each) were used (Buzsaki16-CM16LP, Neuronexus). The probe was connected to an headstage preamplifier (µPA16, Multichannel Systems, Germany) through an Omnetics connector and amplified by a portable 32 channels acquisition amplifier (ME32-FAI-μPA, MultiChannel Systems, Germany) at 20 kHz sampling rate.

### Optogenetic stimulation

During the recording session, the implanted cannula (zirconia cannula, Prizmatix. 1.25mm external diameter. 0.5mm diameter of the core optic fiber) is connected to an optic fiber (NA=0.63, 0.5mm core, Prizmatix). Blue light (16 mW/mm2) is generated by a LED controller (Prizmatix; UHP-T-LED 460) and stimulation frequency was 125Hz. Three additional frequencies were tested; however, they were not analyzed because too few usable trials were available.

### Behavior

Mice were water restricted at least one week after surgeries, after the animals have fully recovered from surgeries. During water restriction mice were given access to water 30 minutes per day ad libitum at the same hour. Body weight and general health status were checked daily. If the mouse lost more than 20% of their original weight, the procedure was stopped, and the mouse removed from the experimental group.

The task: two visual cues (0.5s each. Cue 1: a triangle, Cue 2: a square) were displayed in front of the animal with an interval of 1 sec. After the second cue a drop of water is delivered for 150 msec after a random delay (0.4 s or 0.9 s with a random distribution) corresponding to a variable delay. The last day of the experiment, the random delay was replaced by a fixed delay (0.5s). The volume of water delivered during a session was around 3mL. Throughout all sessions, mice were allowed to run freely on an unmotorized wheel, and running speed was continuously monitored and recorded. Mice were trained for 11 consecutive days (2 sessions a day of 30 minutes each, in the light phase). The duration of the training was variable as it depended on the performance of the individual mouse. The spout was gradually moved away from the mouth of the mouse until it reached the final position. Training was considered finished when the mouse was able to lick reliably from the water spout in the final position for 3 consecutive training sessions. The last two days of the protocol were reserved for experimental sessions, starting with the variable delay condition on the first day, followed by the fixed delay condition on the final day. Each experimental day included a control sessions starting with 60 trials with no task, only recordings and 60 trials with the behavioral task. Stimulation session followed with 30 task trials without stimulation followed by 30 task trials with photostimulation.

### Mapping cerebello-prefrontal functional connections in anesthetized mice

Mice were anesthetized as for *surgeries*. A frontal craniotomy was performed as for *surgeries*, while a wider craniotomy above the right cerebellum was performed as a rectangle of 3mm in width and 2mm in height to expose the entire right posterior lobe. Craniotomies were kept moist with NaCl 0.9% warmed at 37 °C. An acute probe (16 channels, 4×4 tetrodes, Atlas Neuroengineering) was lowered in the left PrL (same coordinates as above) while optogenetic stimulation (optic fiber, 0.5mm core diameter, 0.63 NA) was performed on the cerebellar cortex and manually moved to the different stimulating areas. Six different areas of the cerebellar cortex were stimulated: lobule VI, lobule VII, CrusI and CrusII medial (paravermis) and CrusI and CrusII lateral (hemispheres). Phostimulation was steady (1s) or as pulses (4 pulses of 0.5s every 1s, 4s in total). 25 Recordings for each protocol were done on each mouse at each location. At the end of the experiment, mice were euthanized with Euthasol (400 mg/kg) while still under anesthesia. Data were acquired as described above (see *Recordings*). The 16 channels were averaged and bandpass filtered to 0.1 - 60Hz. All the recordings from the same mouse with the same condition were averaged together and the negative peak from the stimulation window and a baseline of same duration (1 s or 4s for steady and pulses protocol, respectively) were determined.

### Analysis of Local Field Potential

Analyses were performed using Python Libraries (Numpy, Scipy, Elephant, Neo) on recordings. The local field potential (LFP) was extracted from the electrophysiological recordings. Four bands of biologically relevant frequencies of oscillation were considered in our analyses: delta (1.5 – 4Hz), theta (4 – 10Hz), beta (10 – 30Hz), and gamma (30 – 80Hz)(Buzsáki and Draguhn, 2004). Time-frequency analyses were performed using continuous wavelet transform (CWT)(KRONLAND-MARTINET et al., 1987; Roux et al., 2007) to extract the amplitude of each frequency band during the task. Amplitudes were averaged across channels and recordings of the same sessions for each mouse and averaged across mice. Power spectral density was calculated by Welch’s method. The signal of each electrode was individually downsampled to 1kHz, segment length of 1s with an overlap of 0.5s were used. Frequencies from 1 to 80Hz of the resulting power spectrum were kept and normalized to the total power. Then the total power of each band of interest was averaged between trials of the same mouse and then across all mice.

### Spike sorting and PCA

Spiking activity was extracted using the SpikeInterface toolbox(Buccino et al., 2020). For each session, recordings from all trials were concatenated before preprocessing. Signals were bandpass filtered between 300 and 6000 Hz to suppress low-frequency components, including LFP activity, and to remove high-frequency noise. To reduce noise shared across channels, the median signal was subtracted from each channel. Putative units were then identified with the Kilosort 2.5 algorithm. After automated sorting, units were curated manually to discard noise clusters and multiunit activity, and only well-isolated single units were retained for further analysis. The instantaneous firing rate for each neuron was obtained by convolving its spike train with a Gaussian kernel with a sigma of 200 ms.

To identify latent patterns of activity across the population of recorded units, we performed a principal component analysis (PCA) using only the first 4 s of each trial, during which all behavioral events occurred. Each neuron’s mean firing rate was treated as a variable, and each time point served as an observation. The first three principal components were used as templates to identify clusters of units exhibiting similar activity patterns. For each neuron, we computed the Pearson correlation between its firing rate and each principal component. Each neuron was then assigned to the component with which it had the strongest positive or negative correlation, as long as the correlation coefficient was greater than 0.5 and statistically significant (p < 0.05).

### Inter Trial Phase Consistency (ITPC)

The local field potential was extracted from electrophysiological recordings and band pass filtered for the four frequency bands defined above. Phase of the signal for each band, recording and channel was extracted using the Hilbert transform and taking the angle as in (Breska and Ivry, 2020). Phase signal was then aligned per trial for each mouse and band and ITPC was computed then average across channel.

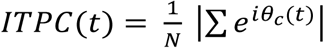

Where (𝑡) is the instantaneous phase of the filtered recording obtain via the Hilbert transform at time t and N is the number of trials. This index measures, at each time point, the degree of phase alignment across trials. Its value is bounded between 0 and 1, with values close to 1 indicating perfect alignment and values close to 0 indicating no alignment.

### Classifier analysis

To assess whether photostimulation altered neuronal activity, we trained a supervised logistic-regression decoder to predict trial identity (“control” vs “photostimulation”) from single-trial firing-rate profiles, evaluated for each 50-ms time bin. For each unit and each trial, spike counts were binned (50 ms bin width) and converted to firing rates. Decoder performance was quantified as the area under the receiver-operating characteristic (auROC) computed separately for each time bin as in (Cohen et al., 2012). Performance estimates were obtained with 2-fold cross-validation, and the reported auROC at each bin is the mean across folds, yielding a temporal auROC profile per neuron. For each condition and time bin, the mean auROC across units was compared against the theoretical chance level of 0.5 using a one-sample t-test. This test evaluated whether decoding accuracy differed significantly from chance. When the resulting p-value was below 0.05, the mean auROC was considered significantly above chance, indicating reliable trial discrimination based on neuronal activity alone.

To examine population structure, auROC time series were embedded via principal component analysis (PCA; 10 components) and time bins were clustered by hierarchical clustering in the reduced PCA space. Four clusters were identified for the variable 400-ms and fixed-delay conditions, and three clusters for the variable 900-ms condition. Neurons whose peak auROC occurred within the cluster corresponding to the reward period were designated as photomodulated.

### Statistics

Comparisons of the slopes of the different metrics used to evaluate anticipation during the delay were performed after a bootstrap procedure (Efron and Tibshirani, 1993). Bootstrapping reduces noise and captures general trends in small datasets by resampling with replacement to generate multiple simulated samples. For each mouse and condition, linear regression was computed on the delay distribution, and the resulting slope values were bootstrapped to estimate their variability and group-level trends. Shuffled distributions, obtained by randomizing data points across time before regression, were used as a baseline for comparison. When comparing two bootstrapped distributions, a Cohen’s d effect size was computed to quantify the magnitude of the difference, with values ≥ 0.8 considered significant.

In addition to slope-based metrics, inter-trial phase consistency (ITPC) was quantified to assess temporal alignment of neural activity across trials when comparing fixed and variable delay conditions. ITPC was computed on data downsampled to 100 Hz and a cumulative sum of the resulting ITPC from reward delivery to reward removal was performed. Statistical differences between conditions were assessed using a Kolmogorov–Smirnov test to determine whether the ITPC distributions differed significantly.

Purkinje-cell stimulation trials were delivered later within each session, at which point baseline levels of several behavioral and neural measures (e.g., licking rate, running speed, and neuronal firing rate) typically exhibited temporal drift. To mitigate this drift and enable valid between-condition comparisons, each metric was mean-centered by subtracting its session- and condition-specific average. This normalization evaluated stimulation effects relative to each animal’s contemporaneous baseline rather than absolute values, thereby reducing confounds arising from intra-session drift.

To evaluate the effect of Purkinje cell photostimulation, differences between stimulated and non-stimulated trials were tested using a repeated-measures two-way ANOVA. For each mouse, the analysis covered the interval from the first possible reward time at 2.4 s to the end of the last possible reward window at 3.05 s. The resulting data were then entered into the ANOVA with time and condition as factors. Effects were considered significant when either the main effect of condition or the interaction between time and condition reached significance.

## Supplementary Information

**Supplementary Figure 1.**
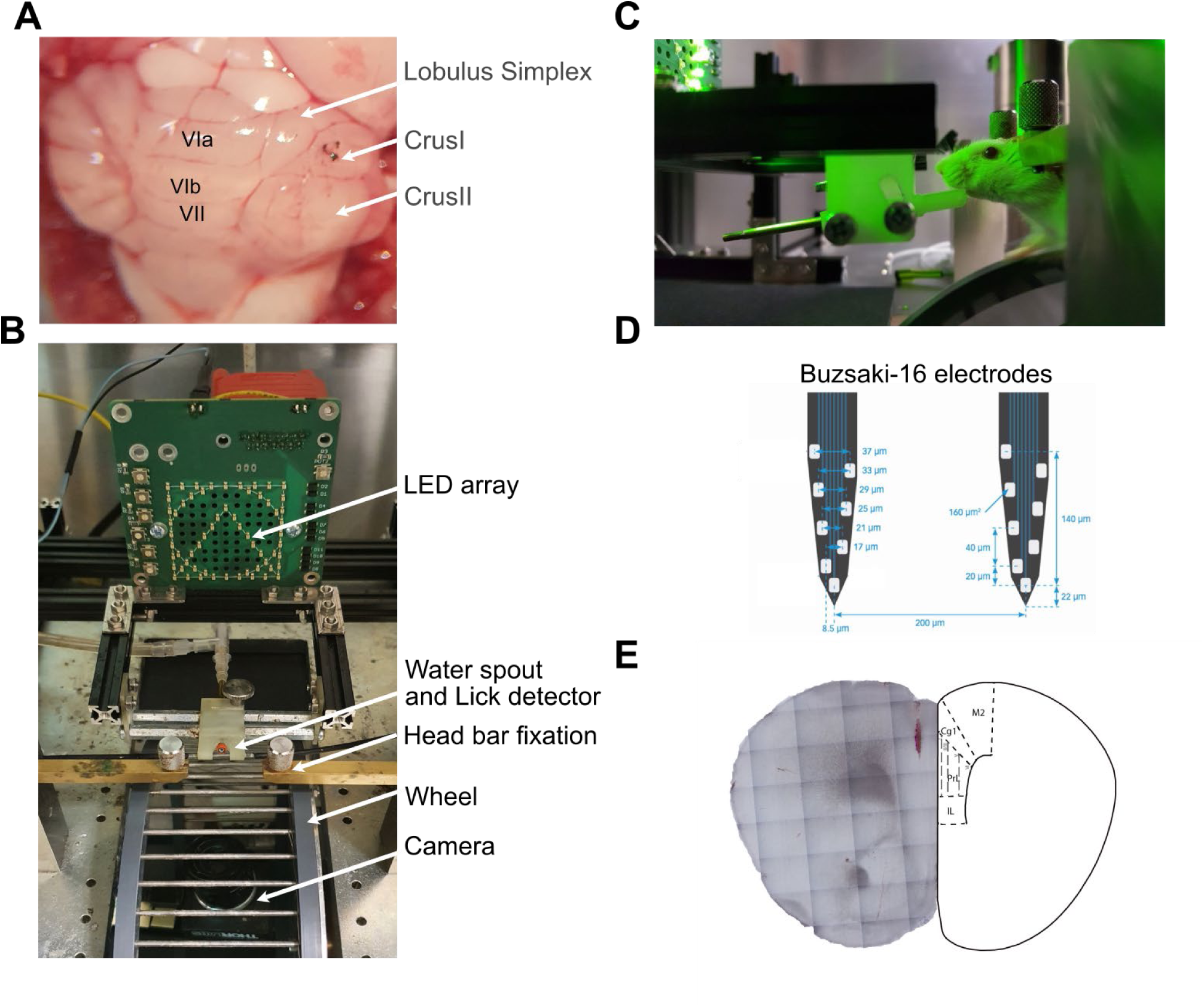
Experimental setup and recording configuration. **A.** Dorsal view of the cerebellar surface. Otogenetic fiber is implantated above Crus I. **B.** Head-fixed behavioral setup with LED visual cue array, water spout with lick detector, head bar fixation, wheel for voluntary running, and behavioral monitoring camera. **C.** Image of a mouse during an experimental session.**D.** Schematic of the Buzsáki-16 silicon probe used for electrophysiological recordings in the medial prefrontal cortex (mPFC), showing electrode spacing and configuration. **E.** Histological verification of electrode placement within the mPFC.

**Supplementary Figure 2.**
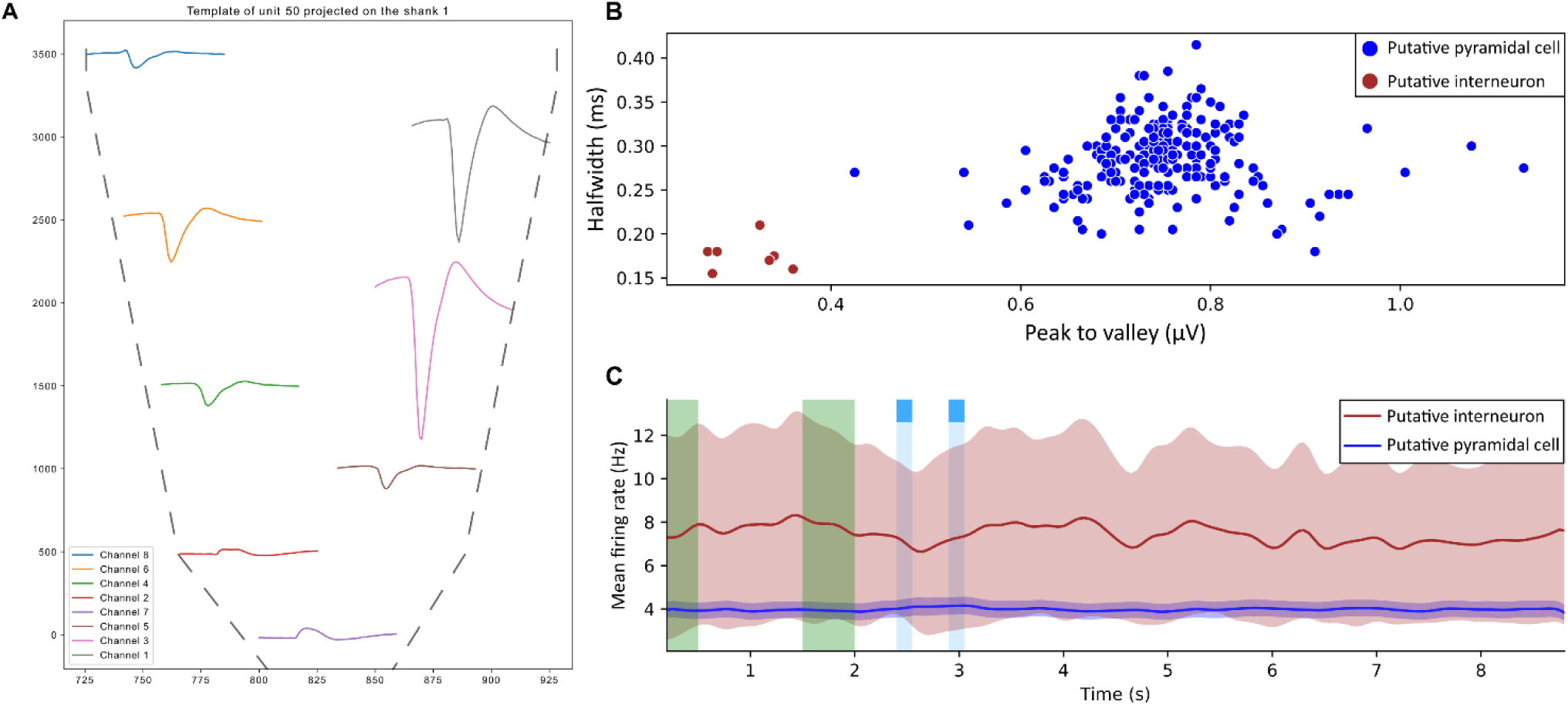
Unit classification. **A.** Example of spike waveform recorded across channels of a Buzsáki-16 probe. **B.** Scatter plot of spike half-widths versus peak-to-valley amplitudes used to classify putative pyramidal neurons (blue) and interneurons (red). **C.** Averaged firing rate of putative pyramidal cells and interneurons aligned to task events. Green and blue shaded areas indicate cue and reward periods, respectively. Shaded regions represent SEM.

**Supplementary Figure 3.**
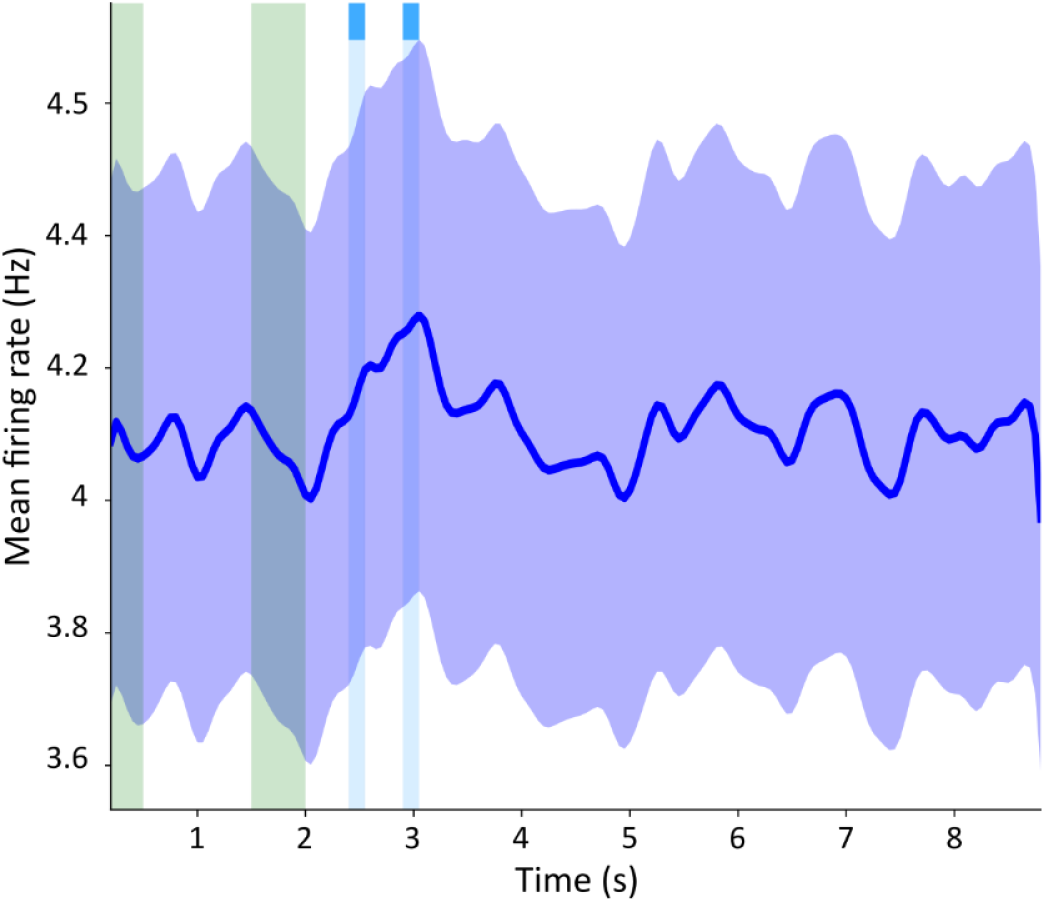
Mean firing rate of mPFC neurons during the timing task. Average firing rate of all recorded units aligned to task events across trials. The green shaded area marks the cue presentation period, and the blue shaded area indicates the reward delivery period. Shaded regions represent SEM.

**Supplementary Figure 4.**
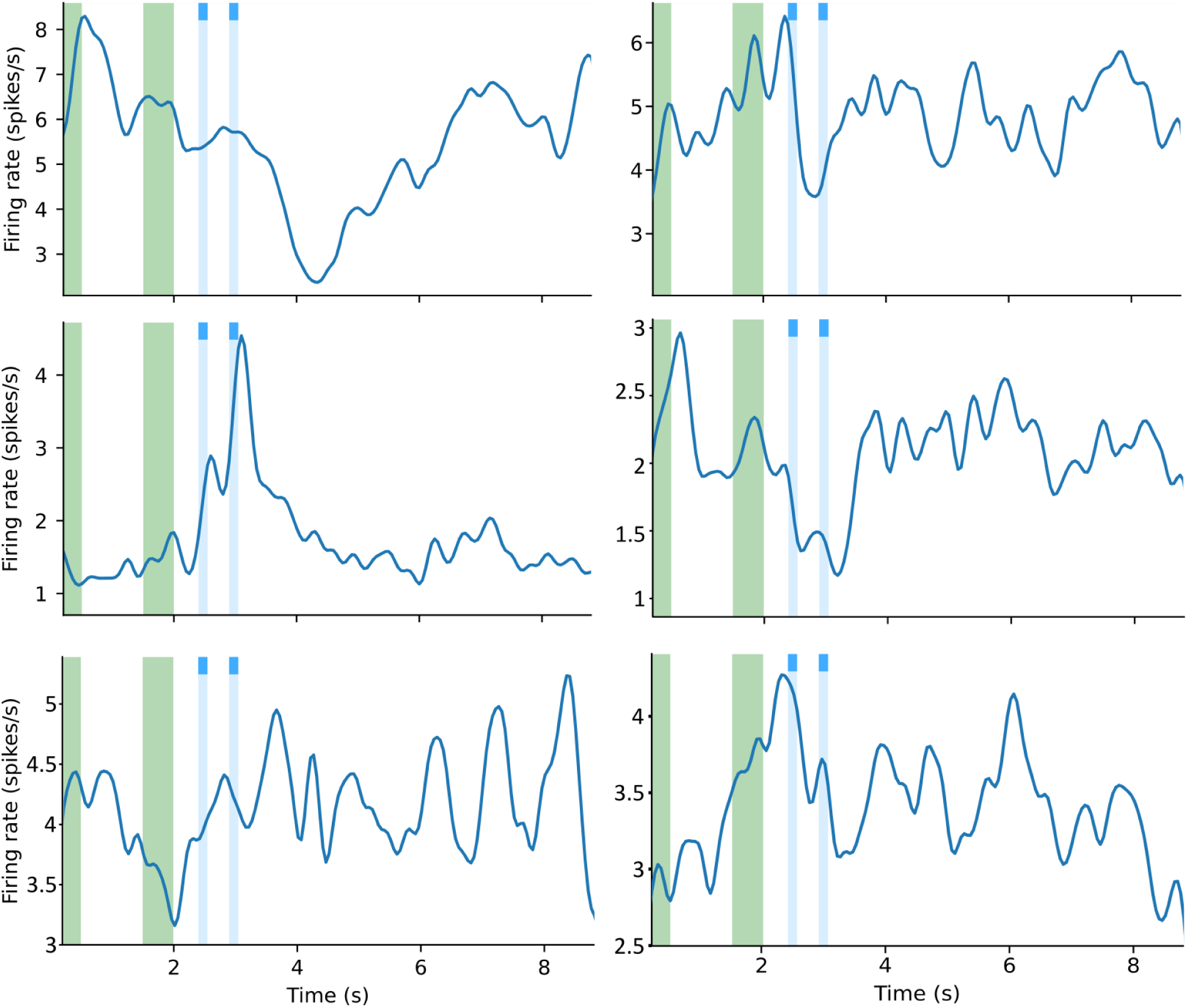
Representative examples of mPFC neuron firing during the timing task. Firing rate profiles of six individual prelimbic neurons aligned to cue and reward events. The green shaded area marks the cue presentation, and the blue shaded area indicates the timing of reward delivery.

**Supplementary Figure 5.**
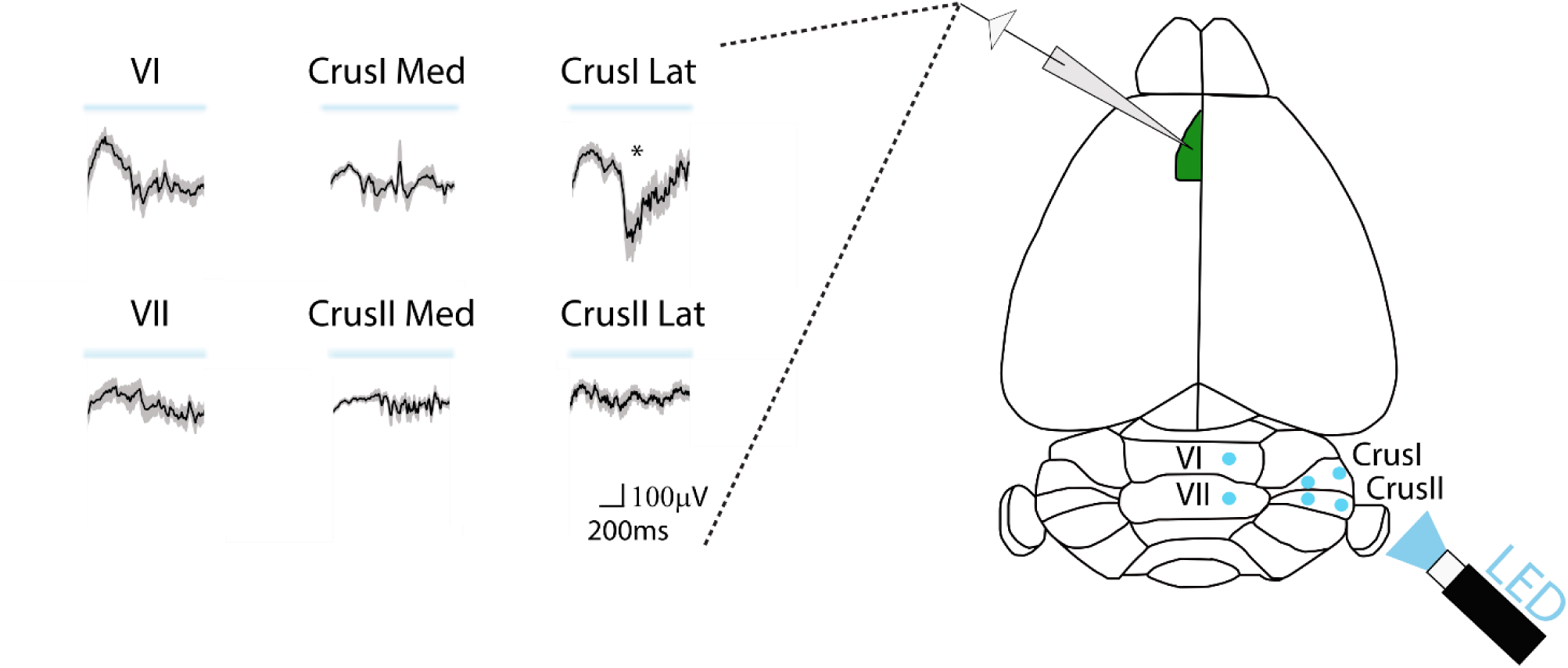
LFP in mPFC following cerebellar photostimulation in anesthetized animals. ***Left,*** Mean Local field potential responses recorded in different cerebellar lobules during optogenetic stimulation (N=5 mice). Traces represent averaged LFPs (± SEM) from lobules VI, VII, Crus I, and Crus II (medial and lateral). ***Right,*** Schematic dorsal view of cerebellar stimulation and recording sites. Blue circles indicate optically stimulated regions; the asterisk marks the site showing the largest evoked response.

**Supplementary Figure 6.**
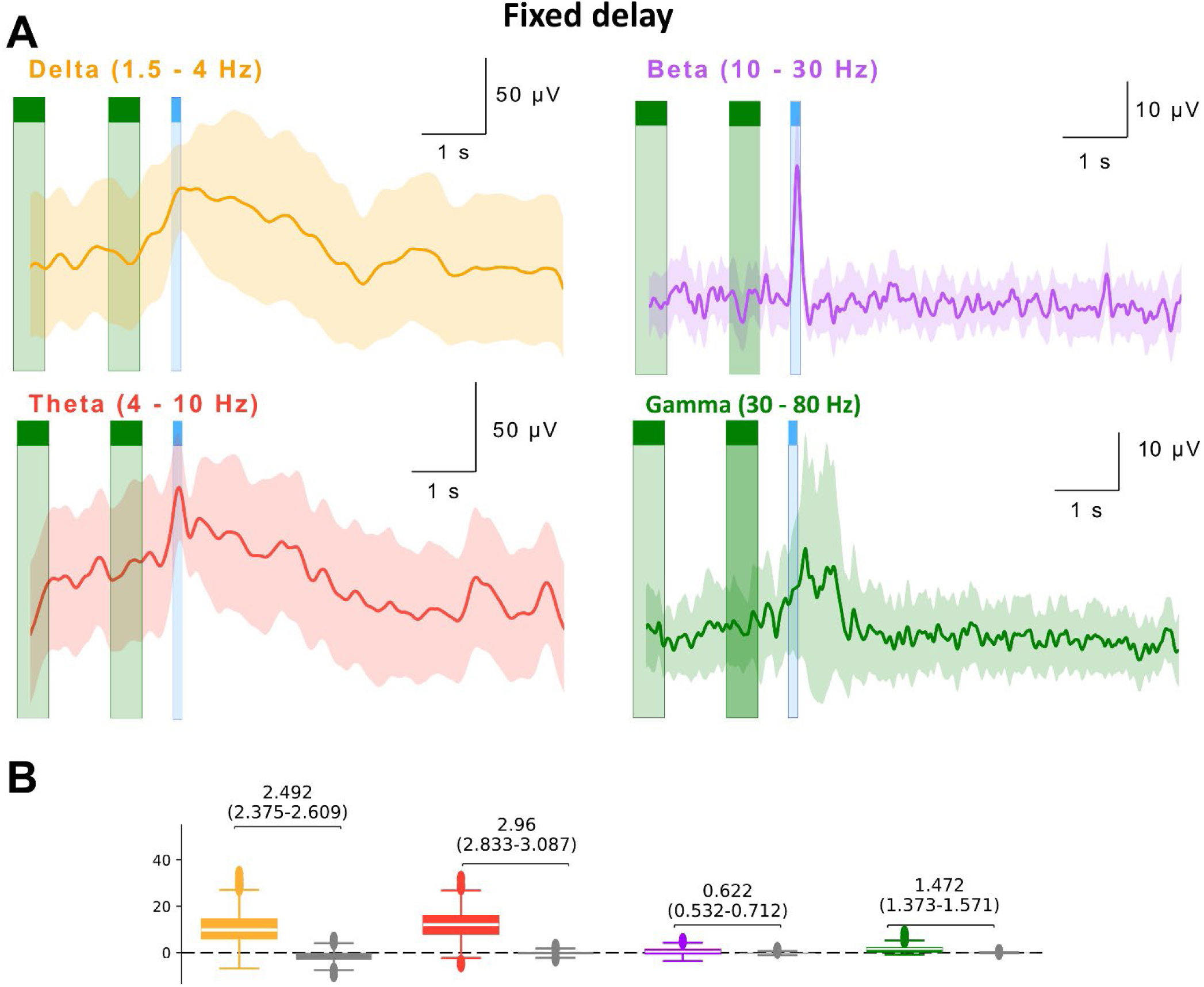
LFP in mPFC during fixed delays., **A,** Time course of averaged band amplitudes across mice (N=9). **B,** bootstrapped distributions of the slope of the averaged amplitudes computed from the anticipatory period for each band, compared with a permutation-based null distribution generated by time-resampling within trials (N=9). Reported values correspond to Cohen’s d effect size, quantifying the strength of the anticipatory increase in LFP amplitude relative to this null distribution.

**Supplementary Figure 7.**
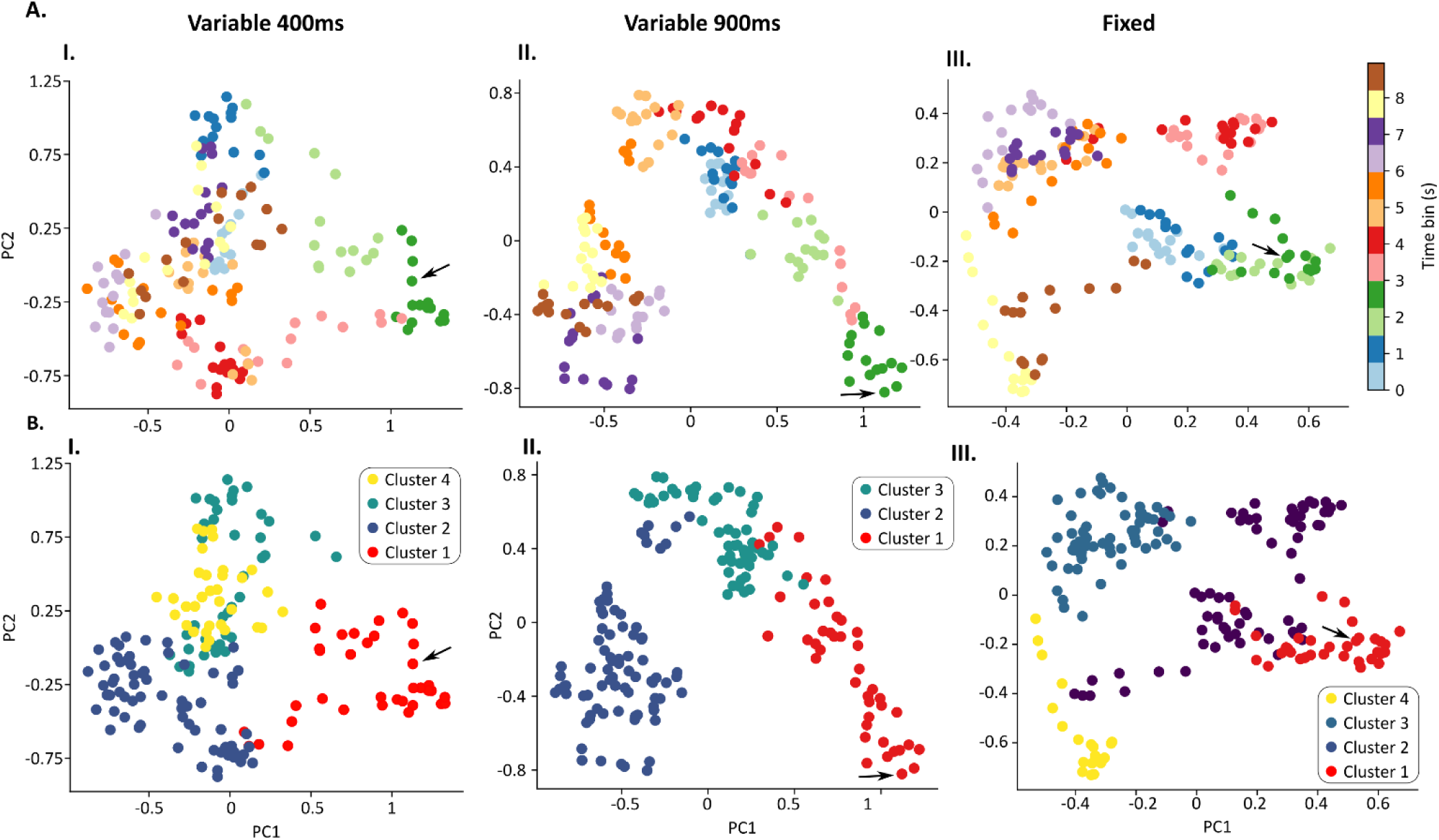
Principal component analysis of auROC and epoch clusters. **A,** principal component analysis (PCA) applied to the population auROC values, where each dot represents a single time bin. Dots are color-coded according to time within the trial. The black arrow marks the time of the first possible reward (2.4 s). The PCA reveals a continuous trajectory across time, with a clear excursion around the reward period. **B**, hierarchical clustering performed on PCA data identified distinct temporal clusters corresponding to different trial epochs. The time around reward formed a separate cluster (shown in red).

